# Quantitative comparison of manuka and clover honey proteomes with royal jelly

**DOI:** 10.1101/2022.07.30.502129

**Authors:** B.W. Paget, T. Kleffmann, K.E. Whiteman, M.F. Thomas, C.D. McMahon

## Abstract

Royal jelly and honey are two substances produced successively by the worker bee caste. Sequential window acquisition of all theoretical fragment-ion spectra mass spectrometry (SWATH-MS) was used to compare protein quantities of bee origin in manuka and clover honey to royal jelly. Two analysis techniques identified 75 proteins in total. Peptide intensity was directly compared for a subset of 32 proteins that were identified with high confidence, and the relative changes in protein abundance were compared between each honey and royal jelly. A total of 16 proteins were found only in honey, 9 of which were common to manuka and clover honeys, and 7 found only in manuka honey. None of the identified proteins were unique to clover honey, while four proteins were unique to royal jelly. Major Royal Jelly Proteins (MRJPs) had similar profiles in both honeys, except MRJP6, which was significantly more abundant in clover honey.

Quantitative data suggests changes of the protein composition as bees develop through the worker caste, with subtle changes in MRJP amounts, and extensive changes in proteins with enzymatic function. Metabolism enzymes were more abundant in honey than in royal jelly, where the trend revealed a potential catalytic role for two uncharacterised proteins (MRJP6 and LOC408608). Differences in the proteomes of the two types of honey probably reflects differences in the type of nectar foraged.

## Introduction

The worker bee caste performs hive functions in a temporal manner, which includes secretion of royal jelly by nurse bees, and production of honey via nectar foraging bees. After emergence from the larval stage, nurse bees begin to produce royal jelly as a secretion from the hypopharyngeal and mandibular glands for the development and nutrition of larvae and adult queens. Protein constitutes 10-15% (w/w) of this secretion, of which the MRJPs and, in particular, MRJP1-5 comprise up to 90% of the protein content [1-5].

The principal task of the forager bee is to collect nectar, the precursor material of honey. Honey is a viscous sweet substance produced in and secreted from the hypopharyngeal glands of forager bees and then stored in the comb of beehives to sustain the energy needs of the hive. It consists mainly of fructose, glucose, and water, with a much lower protein content (mean 0.17% (w/w), range 0.06 – 0.78%) compared with royal jelly [6, 7]. The secretions become mixed with nectar during collection resulting in the mass of protein being much lower than in royal jelly.

Although the protein component of royal jelly and honey is produced mainly by the hypopharyngeal glands, little is known about the comparative differences in protein composition of the two substances. This arises in part due to the difficulty in comprehensively characterising the protein component of honey and has potentially led to assumptions about the similarity of the protein composition of honey and royal jelly.

Counter to any assumptions, collected nectar is stored in the honey crop, which secretes mono- and polysaccharide digestive enzymes into the nectar bolus [8], and recent research shows that expression of some of the major royal jelly proteins is different in the heads of nurse and forager bees [9]. Additionally, the two products serve completely different functions in the beehive. Taken together, these observations suggest differences should exist in the relative abundance of protein types between honey and royal jelly.

We employed a label-free quantitation technique termed SWATH-MS to investigate and measure potential differences between two honey varieties (manuka and clover honey) and royal jelly produced by *A. mellifera*. Manuka honey is produced from nectar collected from flowers of the *Leptospermum scoparium* plant that is indigenous to New Zealand. Manuka honey has broad spectrum antibacterial activity [10] that is attributed to the presence of methylglyoxal (MGO) [11-13]. Medical grade manuka honey has greater than 263 mg/kg of MGO (10+ on the UMF grading scale), and honey with concentrations above 572 mg/kg (UMF 16+) are of a premium grade [13, 14]. In contrast, clover honey is produced from nectar collected from *Trifolium* species, which are grown in temperate regions worldwide to fix nitrogen in soil to aid growth of pasture for livestock production [15]. The honey has trace amounts of MGO and has limited antibacterial actions compared to that of manuka honey [16].

## Materials and Methods

### Honey and royal jelly samples

Six different clover honeys were purchased from public retail outlets. Pure clover honeys were from Lorimer’s Honey (Hamilton, New Zealand), Pams (Foodstuffs, Auckland, New Zealand) and Airborne Honey (Leeston, New Zealand). Clover blend honeys were from Arataki Honey (Havelock North, New Zealand), Airborne and Mother Earth (Hamilton, New Zealand).

Twelve manuka honeys were sourced from Mother Earth, Red Seal (Auckland, New Zealand), Settlers Honey (Whanganui, New Zealand), and Watson & Son (Masterton, New Zealand).Eight royal jelly samples were analysed. Four fresh royal jelly samples were purchased from Beaut Bees (Auckland, New Zealand), which were harvested on four different dates in 2018. A single fresh royal jelly sample was purchased from Happy Valley Honey (Auckland, New Zealand). ManukaMed (Masterton, New Zealand) provided royal jelly harvested in the summer of 2016/2017 and frozen at - 20 °C until analysis. Royal jelly in capsule form was purchased online from Comvita (Paengaroa, New Zealand) and Manuka Health (Te Awamutu, New Zealand).

### Protein extraction

Honey protein extracts were prepared by diluting 2 g of honey in 2.6 mL phosphate buffered saline (PBS), pH 7.4. Homogenised samples were then dialysed against 300 mL PBS at 4 °C with gentle stirring, using 10 kDa molecular weight cut-off SnakeSkin dialysis tubing (Thermo Fisher Scientific). Buffer was changed after 3 h and 6 h, and dialysis continued for a further 16 h. Aliquots of 1 mL were centrifuged 18,000 × g for 5 min and the supernatant transferred to a new tube. Royal jelly protein extracts were prepared by dissolving 80 mg in 4 mL PBS then dialysing as described above. Extracts from capsules were prepared by dissolving 540 mg of capsule material in 4 mL PBS. Insoluble capsule material was removed from the solution by brief centrifugation prior to dialysis.

The concentration of protein extracts was determined by BCA assay kit with BSA standards (Merck).

### Mass spectrometry sample preparation

All samples were centrifuged at 16,000 × g for 30 min and 400 µL supernatant was loaded on a 0.5 mL centrifugal ultrafiltration unit with 3 kDa cut-off membrane (Amicon, Merck) and concentrated down to 20 µL. The retentate was reconstituted in 200 µL of 15% acetonitrile (ACN) and 5 mM tris(2-carboxyethyl)phosphine in 200 mM aqueous triethylammonium bicarbonate (TEAB) for denaturing and reduction of disulphide bonds. After 20 min incubation at room temperature the volume was reduced by centrifugation to 20 µL and then reconstituted in 200 µL of 15% ACN and 40 mM iodoacetamide in 200 mM aqueous TEAB for alkylation of free thiol groups. After 15 min incubation in the dark at room temperature the samples were buffer exchange three times to 200 µL 15% ACN in 200 mM aqueous TEAB. After the last buffer exchange, a small aliquot of each sample was used for protein measurement using the Bradford assay (Bio-Rad). Samples were then concentrated down to 20 µL and supplemented with 5 µg of sequencing grade trypsin (Promega) in 15% ACN in 200 mM aqueous TEAB. Proteins were digested on filter at 37 °C overnight and boosted with an additional 2 µg of trypsin in the morning. After an incubation at 37 °C for 4 h the samples were dried using a centrifugal vacuum concentrator.

### Shotgun proteomics

To identify the protein component and build a spectral library, an aliquot from each sample of each group (manuka, clover and royal jelly) was taken and pooled as a representative sample of each group. Each of the three pooled samples was analysed in three technical replicates by data-dependent acquisition using a 5600+ Triple Time-Of-Flight (TOF) mass spectrometer coupled to an Eksigent ekspert nanoLC 415 uHPLC system (AB Sciex). Peptides were loaded onto a 75 µm internal diameter silica emitter tip column packed with Luna (Phenomenex) C18 bead material (3.2 µm, 100 Å) on a length of 20 cm. Peptides were separated by a 90 min LC gradient between mobile phase A (1% ACN, 0.1% formic acid in water) and mobile phase B (0.1% formic acid in 90% aqueous ACN) for manuka and royal jelly pooled samples and a 60 min LC gradient for the clover and royal jelly pooled samples. The mass spectrometer was operated in data-dependent acquisition (DDA) mode using a mass range of 400-1300 m/z for precursor ion measurement and 100-1600 m/z for fragment ion measurement. The top 15 precursor ion signals per cycle were used for collision-induced dissociation (CID) fragment ion measurements at rolling collision energy. Two repeat measurements of each precursor were allowed during a period of 120 s.

### SWATH-MS

Each individual sample was analysed in technical triplicates by data-independent acquisition (DIA) using SWATH-MS for protein quantification. The same LC-methods were used as described for the respective DDA analysis to maintain retention time alignment of DIA spectra with the library. For DIA, 33 consecutive fragment ion spectra with variable m/z isolation window sizes were acquired over a mass range of 315-1250 m/z and an ion accumulation time of 100 ms per spectrum resulting in a total cycle time of 3.3 s. The window sizes were calculated based on the precursor ion densities within the different m/z regions of a representative DDA analysis using the SWATH Variable Window Calculator application (AB Sciex).

### Data analysis and library build

For identification of proteins within groups, data from the DIA analyses were processed using DIA-Umpire version 2.0 [17]. Raw files from manuka and royal jelly 90 min LC runs, and clover 60 min LC runs were converted with ABSCIEX MS Converter (AB Sciex) and msconvert [18]. The files were processed with DIA-Umpire signal extraction module and converted to mzXML format with msconvert. Output files were searched using X!Tandem (2013.06.15.1) with a reference database consisting of all *A. mellifera* entries in the NCBI RefSeq collection (comprising 23,491 sequence entries, downloaded from the NCBI site (https://www.ncbi.nlm.nih.gov/) on 20/04/2021) and reverse decoys. Search parameters included trypsin as the cleavage enzyme, with a maximum of two missed cleavages, 50 ppm precursor mass error, and modifications to cysteine and methionine residues. Search outputs were scored and combined with PeptideProphet [19] and ProteinProphet [20]. The ProteinProphet results for each group were processed through the DIA-Umpire quantification module. The protein false discovery rate (FDR) was set at 1% and filtered with a target-decoy method based on protein probability values. Protein abundance was estimated by MS1 peptide intensity-based absolute quantification (iBAQ). Only data were considered where all three technical replicates of at least two biological replicates identified a protein.

For comparative quantification of proteins, raw data of the DDA analyses were processed and searched against the *A. mellifera* reference database using ProteinPilot (version 4.5, AB Sciex). Search parameters included trypsin as the cleavage enzyme, biological modifications and single amino acid exchanges were allowed as dynamic modifications, and FDR analysis was enabled. The resulting group file was loaded into the SWATH Acquisition MicroApp 2.0 integrated into PeakView (version 2.2, AB Sciex) to build spectral libraries. For accurate retention time alignment and peak matching, a time window of 12 min and a mass accuracy of 50 ppm were allowed. Peak intensities were then extracted from DIA data using the 6 strongest fragment ions from each of the 10 strongest precursors, at a peptide confidence of ≥99% and FDR threshold for peak matching of ≤1%. Shared peptides were excluded. The extracted area under the curve values were imported into MarkerView (version 1.2, AB Sciex) for data normalisation (based on the total sum of peak intensities).

### Statistical analysis

Data were imported into the statistical computing software R [21] via RStudio (version 1.3.1073, Giant Goldenrod) [22]. Venn diagrams were constructed using the package ‘VennDiagram’ [23]. Normalised label-free quantitation data were tested for variance and normality using Fisher’s test and Shapiro-Wilk test, respectively. A non-parametric bootstrap method [24] was used to calculate 95% confidence intervals for biological replicates and p-values for the difference of means, using 10,000 replicates. Graphs were rendered using the package ‘ggplot2’ [25].

## Results

Signal extraction from DIA data using DIA-Umpire and the X!tandem search engine identified 34 different *A. mellifera* proteins across manuka honey, clover honey and royal jelly sample groups (Fig 1A). A total of 10 proteins were common to all three sample groups. These constituted MRJPs 1-5, MRJP7, an isoform of MRJP5, glucose oxidase, a glucosylceramidase-like isoform, and defensin-1. A total of 16 proteins were found only in honey, 9 of which were common to manuka and clover honeys, and 7 found only in manuka honey. None of the identified proteins were unique to clover honey. Four proteins were unique to royal jelly, including apolipophorin-III-like protein, 60S ribosomal protein, lysozyme, and uncharacterised protein LOC551098. Four proteins were common to royal jelly and manuka, but not clover honey. These were MRJP9, icarapin-like protein, MRJP2 isoform X1, and uncharacterized protein LOC413627. S1 Table presents data used to compile Fig 1.

**Fig 1.**
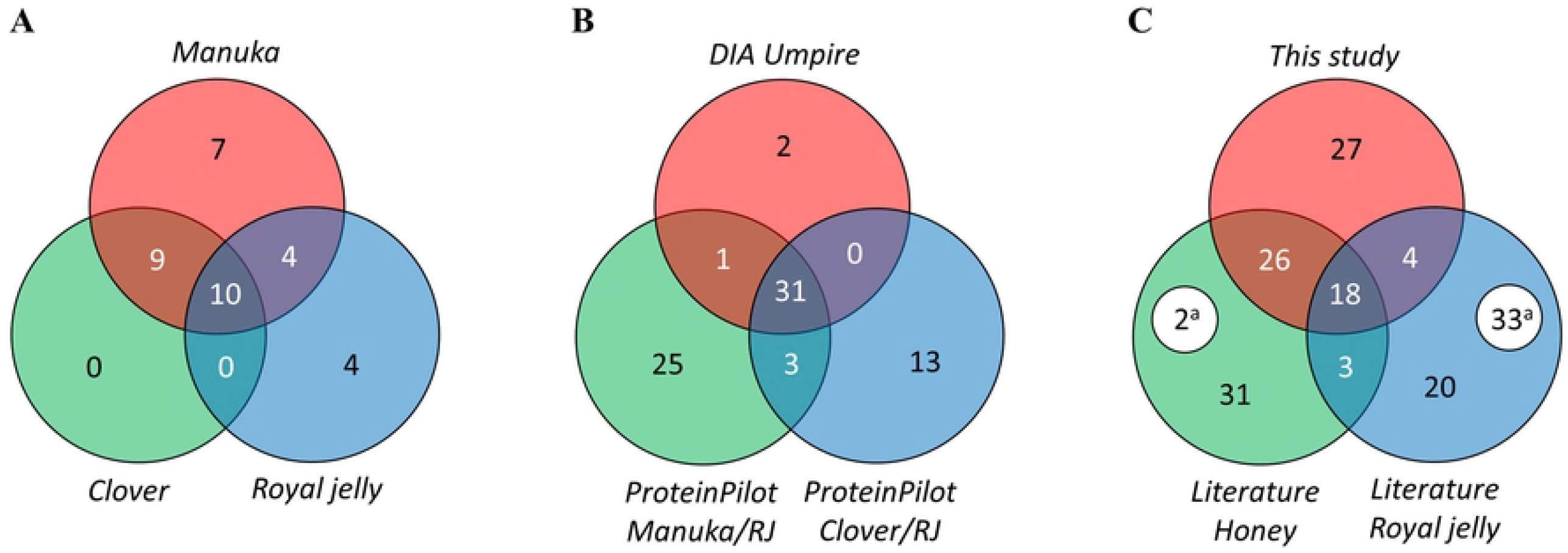
Protein identifications visualised by Venn diagram. (A) Proteins identified in DIA-Umpire by sample group. (B) Proteins identified in DIA-Umpire and the ProteinPilot comparisons of manuka with royal jelly and clover with royal jelly. (C) Proteins identified in this study and proteins identified in the literature in honey and royal jelly. ^a^ proteins removed from the NCBI RefSeq database.

In this study there were 75 proteins identified in total (Fig 1B). The ProteinPilot software using the SWATH-MS workflow identified 73 proteins across all samples, encompassing 32 out of 34 proteins identified by the DIA-Umpire/ X!tandem workflow. The proteins not identified by ProteinPilot were from royal jelly and included the 60S ribosomal protein and uncharacterised protein LOC551098. Criteria for inclusion of data from the literature were proteomic analyses of any honey variety, and of royal jelly. NCBI Identical Protein entries were used to consolidate entries from other databases to create a more concise comparison.

Compared to the literature, there were 27 proteins unique to this study (Fig 1C); 25 were identified by the SWATH-MS workflow and two by the DIA-Umpire workflow. There were 18 proteins that had previously been identified in both honey and royal jelly. These were MRJPs 1-7, MRJP9, glucose oxidase, alpha-amylase, alpha-glucosidase, hymenoptaecin, defensin-1, apolipophorin-III-like protein, putative glucosylceramidase 4, carboxypeptidase Q, uncharacterised protein LOC408608, and ferritin heavy polypeptide-like 17. A further 26 proteins from this study were previously found in honey alone, and 4 previously found in royal jelly alone.

In total, 54 proteins were not identified in this study that were previously identified in the literature – 31 from honey alone, 20 from royal jelly alone, and 3 from both. Due to database processing, 35 identifications in the literature no longer exist in the NCBI RefSeq database. The ProteinProphet results and DIA-Umpire quantification module results from the DIA-Umpire workflow are summarised in Table 1. The inclusion of results by DIA-Umpire was strict. Therefore, the error rates reported by ProteinProphet for manuka, clover and royal jelly identifications at the protein level were 0%, 0.09% and 0%, respectively. Error rates at 1% resulted in 52, 38, and 51 identifications, respectively.

**Table 1.**
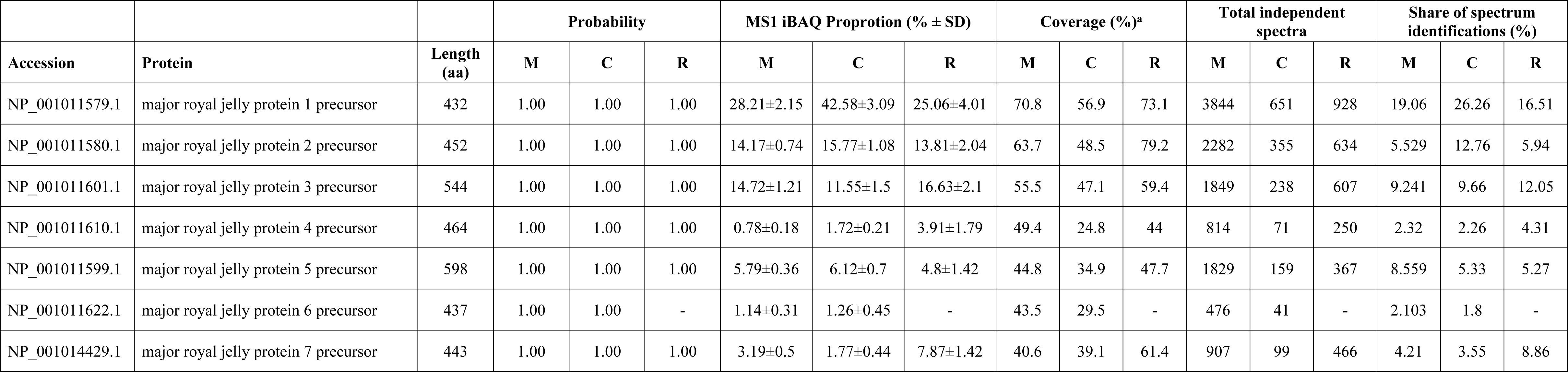
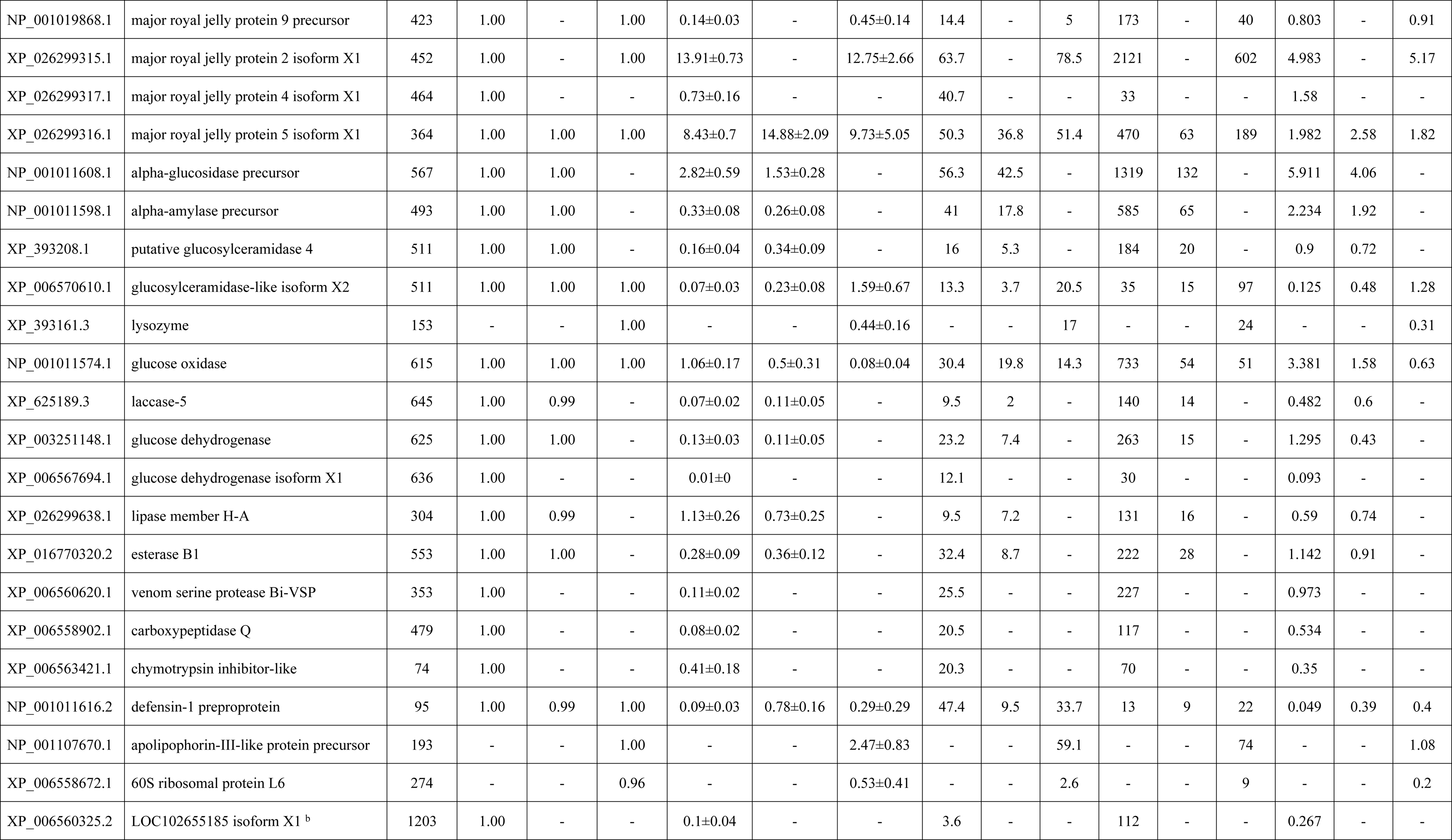
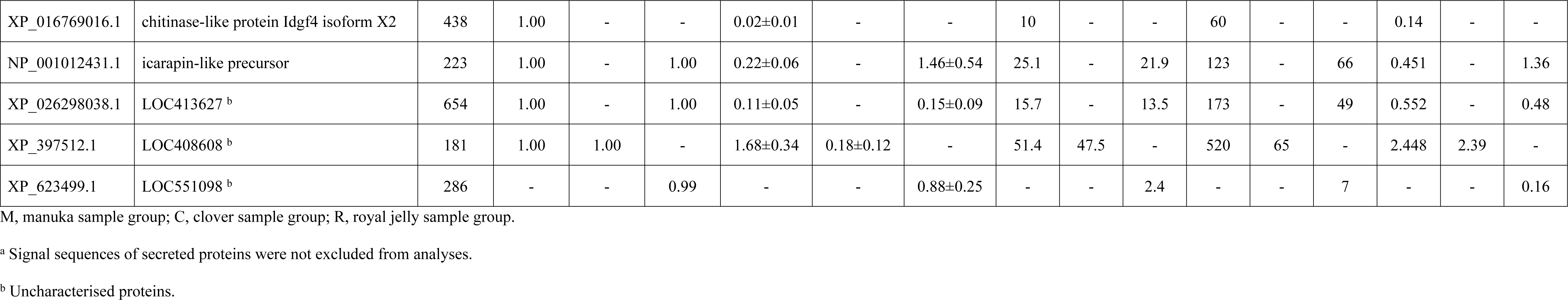
DIA-Umpire and ProteinProphet summary statistics.

The distribution of protein iBAQ values was normal for biological replicates, according to the Shapiro-Wilk test. Across all three groups, MRJP1 was the most abundant protein and had the greatest share of spectrum identifications. In manuka and royal jelly, the proportion of MRJP1 by abundance was 28.21% and 25.06%, respectively. In clover honey, a much higher than expected proportion for MRJP1 was observed at 42.58%. This was likely an artifact of the much lower coverage of MRJP1 than in the other groups, and the absence of MRJP2 isoform X1. The discrepancy was likely compounded by the shorter chromatography run time used for clover samples and highlights the need for caution when comparing semi-absolute quantification data from different experimental methods. The order of abundance of MRJPs varied in each group but accounted for 91.22%, 95.63% and 95.02% of all peptide intensity in manuka, clover and royal jelly, respectively.

Metabolic enzymes made up the remainder of protein abundance for both honeys, many of which were absent from royal jelly. Conversely, the remaining royal jelly proteins were poorly represented or not found in the honey samples. These included glucoslyceramidase-like isoform X2, the lipid transport protein apolipophorin-III-like, the allergen icarapin, and uncharacterised protein LOC551098.

Proteins identified in both analysis workflows (union of Fig 1B) were the focus of comparative quantitative analyses. There were 31 proteins identified by DIA-Umpire workflow and in both SWATH-MS comparisons. Apisimin was also quantified by including a SWATH-MS window between m/z 315-321.4 to monitor the doubly charged peptide precursor TSISVK at m/z 317.69. The SWATH-MS quantitative data was analysed pairwise, comparing royal jelly to both manuka and clover honey separately (Fig 2). Only peptides unique to a protein were included in these analyses to ensure that comparisons between groups were of peptide intensities from the same protein. In doing so, the sum of peak intensities became a less reliable measure of protein abundance. However, in contrast to the DIA-Umpire workflow, the same peptide coverage per protein was considered in each group for the ProteinPilot analysis. Regardless of accuracy, the inclusion of honey protein abundance data as shaded bars in Fig 2 helps to highlight the importance of some fold-change observations.

**Fig 2.**
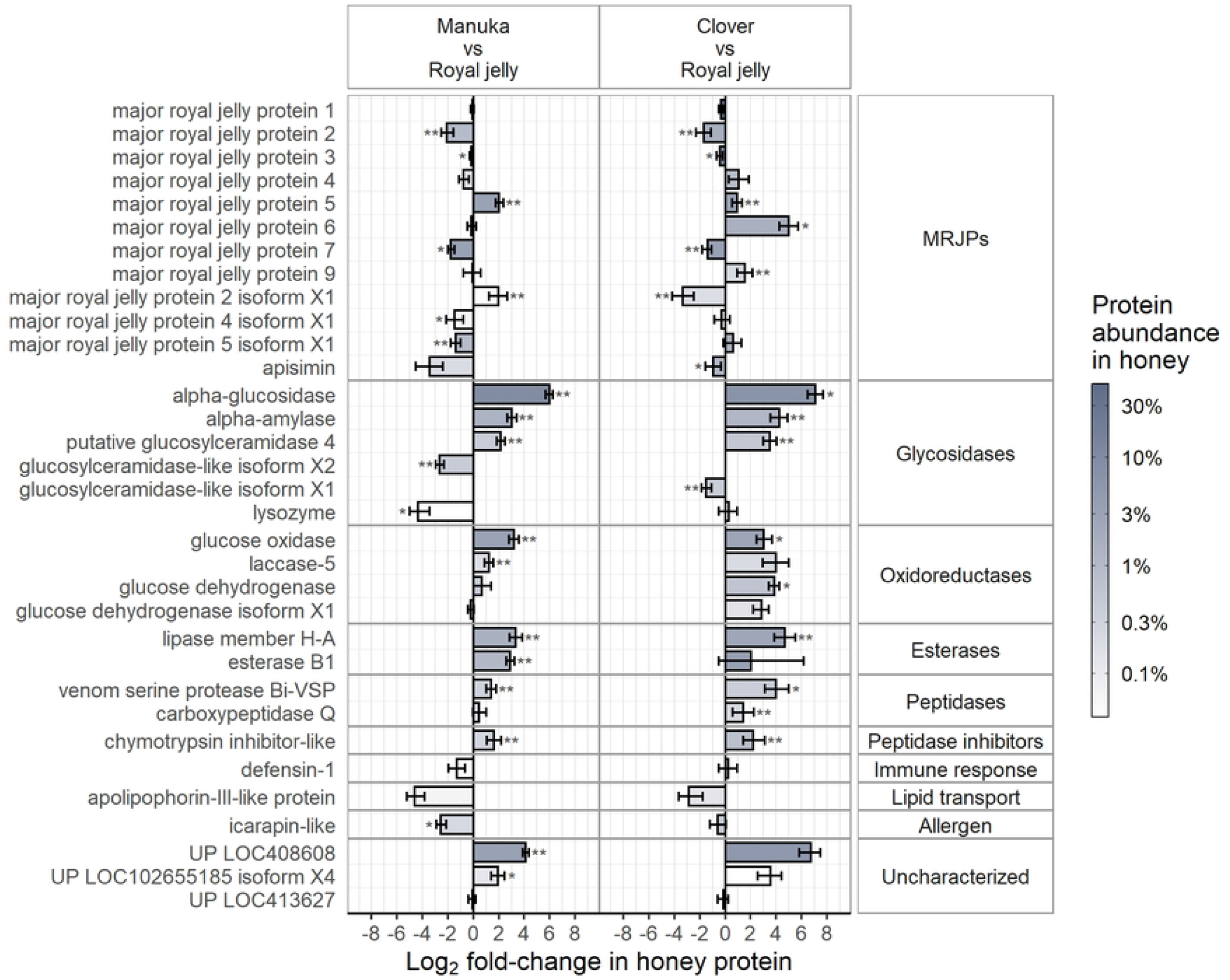
Comparative analyses of manuka and clover honey to royal jelly by SWATH-MS. Quantitative comparisons of proteins identified by both analysis methods, expressed as the log2-fold change in sum of peak intensities for each honey protein compared to the corresponding protein found in royal jelly. Apisimin was added from the SWATH-MS analysis, giving 32 proteins in total. The shading of the bars represents the mean peptide intensity of protein in the honey samples (manuka, n = 12; clover, n = 6). Significant differences are indicated by * *P* < 0.05, ** *P* < 0.01, *** *P* < 0.001.

To look for patterns in relative MRJP abundance, all MRJP proteins were grouped and ordered. Remaining proteins were grouped by enzyme classification or biological role and ordered similarly. MRJP1 and 3 were most abundant in all groups and, therefore, little variation was observed at the Log_2_ scale. As MRJP7 was relatively high in royal jelly, there was 3.39-fold and 2.61-fold less MRJP7 in manuka and clover, respectively. There was significantly more MRJP5 in the honey samples than in royal jelly. MRJP6 was 32.16-fold more abundant in clover than in royal jelly. In contrast, the difference was not significant between manuka and royal jelly. Significantly less MRJP2 was found in both honeys compared to royal jelly. Generally, there were less of the remaining MRJPs in manuka, and more in clover, than in royal jelly.

The non-MRJPs mostly comprised metabolising enzymes. Alpha-glucosidase was most abundant in this group with 63.2-fold and 126-fold more protein in manuka and clover than in royal jelly, respectively. The uncharacterised protein LOC408608 was the next greatest in fold change followed variably by glucose oxidase, alpha-amylase, lipase member H-A, esterase B1, putative glucosylceramidase 4, and glucose dehydrogenase. These enzymes were also consistently found in previous studies in both honey and royal jelly.

A consistent pattern was observed in both manuka and clover honey, wherein the non-MRJPs were present in higher amounts than in royal jelly, except for glucosylceramidase-like isoform X1, apolipophorin-III-like protein, and icarapin-like protein. Additionally, lysozyme and defensin-1 were 20-fold and 2.46-fold lower in manuka honey than in royal jelly.

Venom serine protease Bi-VSP and a chymotrypsin inhibitor-like protein were significantly more abundant in both honeys than in royal jelly, while carboxypeptidase Q was more abundant in clover only.

## Discussion

Until recently, only 17 distinct proteins of bee origin had been identified in various honeys using 1-dimensional [26] and 2-dimensional [27-29] gel electrophoresis techniques coupled with mass spectrometry identification. The low yield of information reflected the intrinsically poor resolution of honey proteins after gel separation, and bias toward selecting bands or spots of clear abundance. In royal jelly, a further 24 proteins had been identified using the same techniques [1, 30-36]. Modern HPLC and high resolution mass spectrometry instruments can circumvent this issue as evidenced by two studies [37, 38] which identified all 18 proteins described by the union of Fig 1C. Other studies using high resolution instruments had identified fewer proteins and focused on post-translational modifications such as glycosylation [39] and phosphorylation [40]. In the present study, shotgun proteomics and relative quantification by label-free SWATH-MS were used on numerous independent samples from manuka honey, clover honey and royal jelly, to provide a comprehensive identification, quantitative analysis, and comparison of these honey varieties with royal jelly. The comparisons were made relative to the extracted protein component, while keeping in mind the large difference in absolute protein content between royal jelly and honey on a mass for mass basis.

### MRJPs

The difference in relative protein expression of MRJPs was generally consistent between the honey types. However, MRJP6 was significantly more abundant in clover honey, while in manuka honey, MRJP6 was comparable to that in royal jelly. Minor differences for MRJPs 4, 9 and the isoforms of 2 and 5 were also observed. The reason for these differences is unclear, although if they also existed at the gene expression level, it may indicate that nectar source has the potential to influence forager MRJP phenotype. Furthermore, the magnitude of the difference in MRJP6 expression in clover honey was similar to non-MRJP proteins with catalytic function, suggesting a possible role in nectar metabolism. Overall, the differences between honey and royal jelly showed some correlation with reported gene expression data of MRJPs from a single bee colony [9], which demonstrated that in the heads of nectar-collecting forager bees, expression of MRJPs 1-4, 7 and 9 was lower, and MRJPs 5 and 6 higher, than in the heads of royal jelly-producing nurse bees.

### Apisimin

Oligomeric MRJP1 consists of a dimerised structure of two MRJP1 molecules bound to two 5 kDa apisimin molecules [41], which is known to polymerise at low pH [42, 43]. Despite using 10 kDa dialysis membrane and relatively high pH buffer, the MRJP1-apisimin dimer was expected to be present in the prepared samples. However, apisimin was not identified in the initial shotgun acquisition, and closer inspection of simulated tryptic digest revealed that a single peptide (TSISVK) would be detected with a doubly charged m/z ratio of 317.69. An additional window was included in the SWATH-MS protocol and apisimin was detected with appreciable intensity. Due to the quaternary structure, the molar ratio of MRJP1 to apisimin was expected to be 1:1. To estimate the ratio in each sample group, the proportional mean intensity was divided by peptide coverage. As a result, there was approximately 26-fold less apisimin than MRJP1 in manuka, 4.9-fold less in clover, and 2.6-fold less in royal jelly. As the amount of MRJP1 was similar for each sample group in the SWATH-MS dataset, the discrepancy in ratios suggests it was not detected in manuka, rather than being absent from the sample. It is likely that apisimin is glycated in manuka honey, and peptide modifications must be known and specified to be detected. While glycations by MGO were included in the ProteinPilot search, the discrepancy suggests more complex glycations, or crosslinks to other peptides might occur, which could also affect trypsin cleavage, and therefore hamper detection in MS experiments.

### Glycosidases, oxidoreductases and esterases

Previous gene expression data has demonstrated upregulation of alpha-glucosidase in the heads of forager bees [44]. Here, glycosidases, oxidoreductases and esterases were greatly upregulated in honey compared to royal jelly. As nectar metabolism is required for production of honey, upregulation across several different enzymes was expected. However, glucosylceramidase-like isoform X2 was more abundant in royal jelly than in both honeys. When compared to the putative glucosylceramidase 4 upregulated in both honeys, similar functional domains are present, with BLAST alignment of 51% identity over 98% of the sequence. It is, therefore, possible that in royal jelly there is a specific role for this glycosidase, having similar catalytic function but distinct substrate specificity. Several additional enzymes were also quantified in the SWATH-MS analysis (S1 Table).

### Immune response and lipid transport

Two peptidases and a single chymotrypsin-like inhibitor were present in both honeys in higher amounts than in royal jelly. Roles in immune defense are likely as the venom serine protease contains a clip domain involved in regulating or localising the immune response [45]. The presence of the chymotrypsin-like inhibitor may help to prevent degradation of bee proteases and other proteins by exogenous proteases.

The amount of defensin-1 was similar in clover honey and royal jelly, in agreement with previous studies [44] showing similar expression of defensin-1 in the heads of nurse and forager bees. It was also demonstrated via immunoblotting that different amounts of mature defensin-1 were present in different varieties of honey, yet it was unclear whether the difference was made up in each honey by unprocessed pro-form defensin-1. The processed state of defensin-1 in manuka honey should not affect the measurements in our experiments due to all quantified peptides belonging to the mature peptide of defensin-1. Therefore, an equivalent amount of defensin-1 should have been observed in manuka and royal jelly. This discrepancy may indicate that actual differences in expression of defensin-1 exist, depending on honey type, as noted for other proteins. As regulation of defensin-1 occurs via the Toll receptor pathway [46], expression should be linked with bacterial burden. In manuka honey, bacterial burden is generally lower due to the presence of MGO, which may indirectly act to reduce Toll receptor signaling and hence expression of defensin-1. Other bactericidal substances in manuka honey [47] would also contribute to this effect. These data support down-regulation of defensin-1 in manuka honey. However, as with apisimin, undetectable glycation PTMs cannot yet be ruled out for the lack of bactericidal activity [44], or peptide abundance in these MS analyses. Additionally, hymenoptaecin was detected in manuka by the SWATH analysis although the sum intensity was nearly 10-fold lower than defensin-1.

Immune-stimulating functions have also been described for insect apolipoproteins [48, 49]. Apolipophorin-III-like protein was less abundant in both honeys, possibly reflecting a more important role in royal jelly, where host defense and lipid availability are essential for larval development.

### Allergens

Icarapin (Api m 10) is one of 13 bee allergens classified in the Allergen Nomenclature database. Peptide coverage of the icarapin-like protein identified in this study was low and did not cover the one amino acid (T140A) difference to the Api m 10 sequence published by the WHO/IUIS Allergen Nomenclature Sub-committee. The official sequence is found in GenBank and Uniprot databases, but not NCBI RefSeq, hence the assignment of icarapin-like.

### Uncharacterised proteins

The uncharacterised protein LOC408608 has been identified several times previously in both honey and royal jelly [30, 35, 37, 38, 40, 50, 51], yet has no clear function assigned. To our knowledge, this is the first time it has been quantitatively compared in the two bee products. The relatively high amount found in both manuka and clover honey is comparable to other metabolising enzymes in these honeys, suggesting a possible catalytic function in honey. NCBI protein-BLAST [52] with non-redundant sequences provided six hits – uncharacterised proteins from five different *Apis* species and one uncharacterised protein from the plant pathogen *Xanthomonas hortorum* – providing no clues to the function. The lack of similarity to known sequences also indicates a species-specific or honey-specific role for this protein.

The uncharacterised protein LOC102655185 is a serine-rich protein with extracellular signal sequence. This was far less abundant in all groups, yet there was at least 4-fold more in honey than in royal jelly. Sequence alignment provided an abundance of hits, many of which aligned to mucins. The uncharacterised protein LOC413627 was similarly abundant in all three groups. Additionally, eight more uncharacterised proteins were detected in the SWATH-MS analysis (Table S1).

## Conclusion

The label-free SWATH-MS approach was able to overcome previous limitations in the quantitative analysis of the honey proteome. Quantitative data suggests expression at the protein level changes as bees develop through the worker caste, with subtle changes in MRJP expression, and extensive changes in proteins with enzymatic function. The latter change has identified at least two proteins that may have a role in nectar metabolism. Subtle differences in the proteomes of the two types of honey probably reflects differences in the type of nectar foraged.

## Supporting information

**S1 Table. Venn diagram data**. Data used to compile Fig 1 Venn diagrams.

